# The Development of Sex Differences in Play in Wild White-Faced Capuchins (*Cebus capucinus*)

**DOI:** 10.1101/2022.07.25.501329

**Authors:** Sasha Lutz Winkler, Susan Emily Perry

## Abstract

Many mammalian species display sex differences in the frequency of play behavior, yet the animal literature includes few longitudinal studies of play, which are important for understanding the developmental timing of sex differences and the evolutionary functions of play. We analyzed social play, solitary play, and grooming using an 18-year dataset on 38 wild white-faced capuchin monkeys (*Cebus capucinus*) followed since infancy. Rates of each behavior were measured as the proportion of point samples taken during focal follows in which the individual engaged in each behavior. To determine sex differences in these rates, we ran a series of generalized linear mixed models, considering both linear and quadratic effects of age, and chose the optimal model for each of the three behavioral outcomes based on information criteria. Rates of both social play and solitary play decreased with age, with the exception of social play in males, which increased in the early juvenile period before decreasing. Male and female capuchins had different temporal patterns of social play, with males playing more than females during most of the juvenile period, but they did not display meaningful sex differences in solitary play rates. Additionally, males and females had different patterns of grooming over the lifespan: males participated in grooming at low rates throughout their lives, while adult females participated in grooming at much higher rates, peaking around age 11 years before declining. We suggest that male and female white-faced capuchins may adopt alternative social bonding strategies, including different developmental timing and different behaviors (social play for males versus grooming for females). Our results were consistent with two functional hypotheses of play, the practice and bonding hypotheses. This study demonstrates that play behavior may be critical for the development of sex-specific social strategies and emphasizes the importance of developmental perspectives on social behaviors.

## INTRODUCTION

Play is both an important and enigmatic behavior in evolutionary science. Although play is costly with regard to time and energy expenditure, it occurs in nearly all young mammals, as well as in other taxa (Burghardt, 2005). As a potential locus of exploration and learning in development, play may have an important role in influencing or predicting adult behavior (Bateson & Martin, 2013; Gopnik et al., 2017). Play is also a domain where sex differences in adult behavior become apparent at a young age in humans and other mammalian species (Barbu et al., 2011; Meaney et al., 1985). While the developmental patterns of play are fairly well studied in humans (Fromberg & Bergen, 2006; Pellegrini & Smith, 1998; Power, 2000), the nonhuman animal literature largely lacks longitudinal data on developmental patterns of play over the lifespan, which are important for understanding the evolutionary forces driving patterns of play across age and sex. The current study investigates such developmental patterns of play in a unique dataset that tracked individuals from several groups of wild white-faced capuchin monkeys (*Cebus capucinus*) over 18 years.

In most mammals, play tends to be more frequent early in life and become less frequent with age (Bateson & Martin, 2013). Play among adults is rare or even nonexistent in some animal species, although some, like humans, continue to play into adulthood (Byers & Walker, 1995). Generally, more precocial species tend to have lower rates of play and a shorter developmental period during which play occurs (Ortega & Bekoff, 1987). This emphasizes the potential role of play in learning. Sexual maturity may mark an important developmental milestone for play; as sexually mature animals increase the time devoted to behaviors that more directly improve reproductive success, like mating and parental care, play appears to decrease in frequency. Even in species that continue to play as adults, rates of play are almost invariably higher in immature animals.

The evolutionary function of play behavior has long puzzled biologists, as the fitness benefits of play are difficult to detect and measure. An additional complexity is that different types of play have likely evolved for different reasons (Burghardt, 2010; Smaldino et al., 2019). Play is often split into two categories: ***social play***, such as rough-and-tumble play, and ***solitary play***, which includes non-social object and locomotor play (Pellegrini & Smith, 2005). In his seminal book on play in rhesus macaques (*Macaca mulatta*), Donald Symons detailed a non-exhaustive list of eighteen different functional hypotheses for the existence of play (Symons, 1978). While some of these hypotheses posit immediate, primarily physical benefits of play, many propose that play in the juvenile period translates to delayed fitness benefits in adulthood (Burghardt, 2010). One such hypothesis is the ***practice hypothesis***. According to the practice hypothesis, play provides practice for skills needed in adulthood; for example, social rough-and-tumble play (i.e., “play fighting”) is seen as practice for skills needed during real fights in adulthood, such as physical agility, tactical skills, and unpredictable counterattacks (Martin & Caro, 1985; Špinka et al., 2001; Symons, 1978).

Another hypothesis for the function of play, particularly for social primates, is the ***bonding hypothesis***. The bonding hypothesis argues that playing with others allows animals to form, test, and learn about social bonds (Maestripieri & Ross, 2004; Poirier & Smith, 1974). These hypotheses are not mutually exclusive and are difficult to disentangle empirically, as acknowledged by many scholars (e.g., Bateson & Martin, 2013).

The proportion of time and energy devoted to different types of play varies across species (Cordoni et al., 2018; Fry, 2005; Palagi, 2006). Within a given species, the frequency of different types of play may vary across developmental stages and across the sexes (Barbu et al., 2011). Together, these sources of variation suggest that different types of play may confer subtly different evolutionary benefits for males and females, for individuals of different ages, or for species with different socioecological pressures. A better understanding of sex and age differences in play may therefore help to shed further light on the evolutionary functions of play, especially when contextualized within the species’ socioecology. Where we have relevant data, we make some predictions, grounded in the natural history of *Cebus capucinus*, about sex differences that might be expected for each hypothesized function of play described here.

### Sex Differences in Play

The presence of sex differences in the development of play could support either or both of the functional hypotheses discussed: the practice and bonding hypotheses. The ***practice hypothesis*** predicts that the sex that has greater need for physical agility and tactical skills in adulthood will have higher rates of social rough-and-tumble play, which consists of modified aggressive behaviors like gentle biting, wrestling, and chasing. Similarly, the practice hypothesis predicts that the sex with a greater need for extractive foraging or other fitness-relevant object manipulation in adulthood will have higher rates of solitary object play as juveniles. Predictions for the developmental timing of sex differences in play are a bit more obscure. One sex might maximize lifetime play by continuing to play over a longer period of development; alternatively, they might maximize play (and its benefits) by playing more earlier in life in order to reach proficiency at a certain skill more quickly, after which they might stop playing or play at reduced rates, having already gained the needed skill.

In contrast, the ***bonding hypothesis*** predicts higher rates of social play in the sex which gains a greater fitness advantage from having strong social bonds. This may decrease over development if the social bonds (and bond-formation skills) become solidified and further play has diminishing returns; however, if play helps to maintain bonds in adulthood that are important for fitness, one would expect social play to continue throughout the lifespan. The bonding hypothesis provides no specific predictions for solitary play. Regardless, both the practice and bonding theories suggest that sex differences in play should mirror adult sex-specific behavioral strategies for reproduction and survival.

Sex differences in social play are common in cross-sectional research. Studies have found higher rates of social play among males than females in a broad range of mammalian species, from rats (*Rattus norvegicus*) to humans (Auger & Olesen, 2009; Fry, 2005; Poole & Fish, 1976). This sex difference has been observed in many Old World monkeys and apes including gorillas (*Gorilla gorilla*; Maestripieri & Ross, 2004); orangutans (*Pongo abelii*; Rijksen, 1978); rhesus macaques (*Macaca mulatta*) (Brown & Dixson, 2000); vervet monkeys (*Cercopithecus aethiops*; Raleigh et al., 1979), as well as some New World monkeys (e.g., spider monkeys, *Ateles geoffroyi*; Rodrigues, 2014). However, this pattern of higher rates of social play in males is by no means ubiquitous, even among primates. For example, studies have found no sex differences in play in common marmosets (*Callithrix jacchus*; Stevenson & Poole, 1982), coppery titi monkeys (*Callicebus cupreus*; Chau et al., 2008), wolves (*Canis lupus*; Cordoni, 2009), or meerkats (*Suricata suricatta*; Sharpe, 2005). The socioecology and mating system of a species likely has an effect on the development of sex differences in play. For example, there is preliminary evidence that monogamous mating systems are correlated with similar rates of play between males and females, for both social and solitary play (Chau et al., 2008). In species that are monogamous (e.g., titi monkeys) or polyandrous (e.g., marmosets), reduced male-male competition may explain the reduced selection pressure for higher rates of rough-and-tumble play in males.

Other studies have indicated that sex differences in play can vary over the course of development. Research on spider monkeys found that males had higher rates of social play than females as juveniles (Rodrigues, 2014), but another study found the opposite pattern for social play in adulthood, such that females played at higher rates than males (Fedigan & Baxter, 1984). Rodrigues (2014) suggested that female spider monkeys may need to continue playing into adulthood because, as the dispersing sex, they have a continuing need for the bonding benefits of social play compared to males. In hyenas (*Crocuta crocuta*), a species in which females are dominant to males and display many male-typical behaviors and hormonal profiles, one study found that immature females had higher rates of social play than immature males, but there was no difference in rates of solitary object play (Pedersen et al., 1990). However, a longitudinal study in hyenas from infancy to adulthood only found sex differences in the interaction of sex and age on the rate of social play, although that study analyzed rates of initiating social play, rather than rates of the overall time spent in social play (Grebe et al., 2019). The authors found that play initiation decreased with age for both sexes, and the rate of decrease with age was steeper for females than males. Studies like these highlight the need for more research that investigates sex differences in play while accounting for important differences between social and solitary play, and for changes over the course of development. Longitudinal data from wild white-faced capuchins (*Cebus capucinus*) in Costa Rica collected by Perry and colleagues (Perry et al., 2012) provides the rare opportunity to analyze such developmental patterns over decades of research.

### Socioecology and Play in White-Face Capuchins

White-faced capuchins are an excellent species for research on play because they have long juvenile periods, they engage in both social and solitary play, and their social behavior is well-studied (Perry, 2012). Despite being so well-studied, to our knowledge no research has been published on the presence or absence of sex differences in play for this species, or about patterns of play over the lifespan. There is minimal evidence of sex differences in play among other capuchin species (subfamily Cebinae). In a small sample of nine captive juvenile tufted capuchins (*Sapajus apella)*, Paukner and Suomi (2008) found that males had higher rates of social play than females, but there was no sex difference in the rates of solitary play. This study was cross-sectional, with data collected over a four-month period, and did not investigate any changes in the sex difference over development. The current study addresses some of these limitations by analyzing rates of social and solitary play in 38 subjects for up to 18 years of life. Additionally, because the current study concerns wild populations, the results may provide a more ecologically valid perspective on the expression and evolution of play behavior. Rates of play are typically higher in captive animal populations, potentially obscuring sex differences that might occur in the wild (Baldwin & Baldwin, 1974; Himmler et al., 2013).

Because the socioecology of a species might affect developmental patterns of play across the sexes, play research in white-faced capuchins benefits from an understanding of their sex-specific behaviors and reproductive strategies. White-faced capuchins live in multi-male, multi-female groups of approximately 5-40 members, although adult males may spend short periods of their lives in all-male bachelor groups (Perry, 2012). They have unusually long juvenile periods and lifespans for small platyrrhine primates, living up to 37 years in the wild and up to 55 years in captivity (Hakeem et al., 1996; Perry, 2012). Females reach sexual maturity around age 5.5-7.5 years, with mean age of first reproduction being 6.22 years, while the minimum age of first reproduction for males is around 7.5 years old (Perry, 2012; Perry et al., 2012). The extended juvenile period appears to be an important time for developing skills such as extractive foraging techniques (Perry, 2009), some of which may be achieved through object play.

Female white-faced capuchins are philopatric, and males disperse from their natal groups and attempt to join or take over other social groups (Perry et al., 2012). At the Lomas Barbudal field site in Costa Rica, where the data for the current study were collected, the average age of first male migration was 7.6 years, while the average age was 4.5 years among white-faced capuchins in another long-term study at Santa Rosa National Park in Costa Rica (Fedigan & Jack, 2012; Perry et al., 2012). As a result of female philopatry, females generally spend their entire lives with kin (related females) while males do not, although males often co-disperse with their brothers or other related males (Perry, 2012). Males often disperse to different social groups multiple times throughout their lives (Perry, 2012). Males may thus have a greater need to develop social skills for forming bonds with unrelated monkeys in new groups -- skills which, according to the bonding hypothesis, could be developed through social play. Play in the juvenile period could be especially important for males if it allows them to form and test strong bonds with other males before co-dispersing, while still in the relatively safe environment of their natal groups. Despite this, Perry and colleagues (2017) found no effect of the rate of social play in male white-faced capuchins on the age of natal dispersal or the time to obtaining their first alpha position, suggesting that high rates of play early in development do not necessarily speed up the timing of important developmental milestones for male capuchin fitness.

Another factor that might influence rates of social play is white-faced capuchins’ high degree of male reproductive skew. Alpha males monopolize mating opportunities and sire about 96% of the infants born to females that are not their daughters or granddaughters (Godoy et al., 2016; Muniz, 2008). Subordinate males provide important support to the alpha male and females by helping defend the group from predators and out-group males, but they usually do not have access to mating opportunities themselves unless they overthrow the alpha male (Perry & Manson, 2008). An adaptive strategy for some subordinate males may be to increase indirect fitness by supporting related males to achieve alpha status. Thus, male reproductive success is generally dependent on the ability to form aggressive alliances with other males (kin or non-kin) and take over new groups of unrelated females by collaboratively defeating resident males, in order to gain access to mating opportunities (Perry 2012). In contrast, female reproductive success is less skewed, and its variance is likely dependent on competition over access to food resources (Perry et al., 2012; Silk, 1993). The high reproductive skew in males compared to females is important as it puts greater selection pressure on males to (1) develop fighting skills and (2) create strong social bonds with other males, to form the basis of aggressive alliances. Furthermore, fighting skill is known to have a greater impact on mortality for male white-faced capuchins than for females, and aggressive coalitions among females are generally less physically injurious and do not have a substantial impact on their mating opportunities (Gros-Louis et al., 2003; Perry, 1996; Perry, 2012). All of these factors suggest that males should have a greater need for social play during the juvenile period compared to females, whether the benefit is increased fighting skills (i.e., the practice hypothesis) or forming bonds with other males (i.e., the bonding hypothesis). Thus, we predicted higher rates of social play among males than females under both hypotheses.

Previous research on white-faced capuchin social relationships also informed our predictions for the bonding hypothesis. Social bonds are important for both male and female reproductive success; for example, bonded females may help one another through alloparenting and coalitionary aid (Perry, 2012). Our consideration of the bonding functions of play must account for other methods that white-faced capuchins use to form and maintain bonds, such as grooming (Manson et al., 1999). Previous research found that adult females tend to bond with one another by grooming and engaging in relaxed affiliative interactions, while adult males tend to maintain bonds through lower levels of these behaviors, as well as resting in contact, and social play; both sexes participate in innovative dyadic bonding rituals at low rates (Perry, 1996, 1998; Perry et al., 2003). We know that in adulthood, female-female grooming bouts are much more frequent than male-female and male-male grooming (Perry, 1996, 1997, 1998). Perry (2009) found no sex differences in the amount of time immature males and females spent alone or in proximity to their mothers, suggesting that they spend similar overall amounts of time socializing. A more detailed look at the time budgets of immature white-faced capuchins (under age six years) at Lomas Barbudal also supports the idea that males and females spend similar amounts of time socializing (females: average 11.9%, SD=2.6%; males: average 12.6%, SD=2.1%) (Lomas Barbudal Monkey Project, unpublished data). Together, this suggests that males and females in this species tend to establish, maintain, and strengthen social bonds through different means as adults. It is unclear whether this pattern emerges early in development, with juvenile females already allocating more time toward grooming and juvenile males engaging in more social play.

In sum, close dyadic bonds are beneficial for both male and female white-faced capuchins, but because males have the added challenges of integrating into new social groups, forming high-stakes aggressive alliances, and overcoming the odds stacked against them by a high reproductive skew, one could argue that social bonds are likely to provide greater marginal fitness benefits for males than females. Alternatively, female white-faced capuchins may rely more heavily on behaviors such as grooming to form bonds; by replacing social play with grooming, they may continue to gain social bonding benefits but not the fighting benefits associated with social play. Studying sex differences in both play and grooming may thus help to elucidate sex differences in the means by which social bonds are formed and maintained throughout the lifespan.

Finally, what of solitary play? In contrast to social play, solitary (e.g., object and locomotor) play is likely to fulfill the same function for male and female white-faced capuchins. If it is true that different types of play can serve different evolutionary functions, one would expect no sex difference in rates of solitary play when males and females have similar needs for the development of foraging and basic locomotor skills. This may depend on the species’ diet and socioecology. Female reproductive success across primates is generally more dependent on food availability than male reproductive success (Silk 1993), and there is some evidence that adult female white-faced capuchins spend more time foraging than adult males (Rose, 1994). However, there is no evidence of sex differences in basic locomotion, other than differences in the need for fighting skills, as discussed previously. Additionally, the closely-related tufted capuchins (*Sapajus apella*), which have similar socioecologies and extractive foraging niches, do not show sex differences in rates of solitary play (Paukner & Suomi, 2008). Thus, we predicted that it was possible but unlikely that white-faced capuchins would display sex differences in solitary play.

### Research Questions and Predictions

The current study analyzed the patterning of play behavior in white-faced capuchins over the lifespan to answer the following research questions: are there sex differences in the rates of social and solitary play, and how do these sex differences change over development? Additionally, we investigated whether there might be evidence for a trade-off in the activity budgets of males and females between grooming and play. In other words, if social play provides important bond-formation benefits, and females play less often than males, might females compensate for the lack of bonding opportunities by increasing rates of other bond-formation behaviors like grooming? Finally, we were interested in whether the observational evidence provides support for any of the functional hypotheses about play—what, if anything, can sex differences in play tell us about play’s ultimate evolutionary functions?

Based on the patterns of development, dispersal, and play in other primate species and the socioecology of white-faced capuchins, we predicted (1) that rates of both social and solitary play would decrease with age, (2) that males would have higher rates of social play when compared to females throughout the lifespan, and (3) that there would be no sex difference in rates of solitary play.

While most research on play in animals is cross-sectional, (comparing static age groups or sex differences), it is critical to conduct longitudinal studies to understand the timing and emergence of sex- and age-specific patterns. We analyzed the behaviors of a cohort of wild white-faced capuchins that were observed over 18 years (Perry et al., 2012), to see when sex differences emerged and whether sex differences in play were consistent with known sex-specific reproductive strategies.

## METHODS

This research was approved by the Animal Research Committee at the University of California, Los Angeles (ARC 1996-122, 2005-084, 2016-022, and associated renewals) and was conducted in accordance with all federal and international laws of the United States and Costa Rica. Researchers adhered to the American Society of Primatologists’ Principles for the Ethical Treatment of Non-Human Primates and Code for Best Practices in Field Primatology. Research followed the US National Research Council’s Guide for the Care and Use of Laboratory Animals, the US Public Health Service’s Policy on Humane Care and Use of Laboratory Animals, and Guide for the Care and Use of Laboratory Animals as applicable.

### Field Site and Subjects

The data for this study were collected from 2002 to 2020, as part of the Lomas Barbudal Monkey Project, a longitudinal study of wild white-faced capuchin monkeys led by Dr. Susan Perry that began in 1990. The field site includes the Reserva Biológica Lomas Barbudal in the Guanacaste province of Costa Rica and surrounding private areas. Subjects (N=38) included 18 males and 20 females. Subjects were born between August 2000 and April 2005, and were regularly monitored with ongoing behavioral observations beginning in 2002 (or at birth for those born after 2002).

The 38 subjects were born into one of four habituated social groups (group names AA, FF, RR, and FL, which was a fission product of AA). Over the course of the longterm study, the three original groups (AA, RR, FF) fissioned, producing seven additional multi-male, multi-female social groups (FL, MK, CU, CE, DI, RF, SP), and one long-term all-male group (LB; see Supporting Information for more details). Some of the males in this study migrated from their natal group to groups that were observed as part of the Lomas Barbudal Monkey Project, while others migrated to unmonitored groups and were thus lost to the research team, aside from rare glimpses when unhabituated groups were encountered. Thus, throughout the course of this study, subjects were in a total of 11 monitored social groups (those named above), the unmonitored group BD, several small male-only groups, and other unmonitored, unhabituated social groups. We computed, for each subject and year of life, the average group size across all point samples for which we had census data. The average group size across those annual averages for each individual was 23.4 (SD = 7.9, range = 2 to 38.7).

Twenty out of 38 subjects were lost to observation before the end of the study. Of the 20 original females, 10 were presumed to have died, with mean age of death 10.66 years (SD = 5.16 years). Of the 18 original males, only one was observed throughout the entire study period, four were known to have emigrated from the study groups, two likely emigrated from the study groups, nine likely died, and two disappeared for unknown reasons. The average age of disappearance or death for males was 5.86 years (SD 3.51). If males disappeared when they were young and were the only group member to disappear at that time, they were presumed to be dead. If older males disappeared simultaneously with another male from their group, they were presumed to have emigrated outside the study area.

### Data Collection

Rates of social play, solitary play, and grooming were calculated using data from point samples taken during focal follows of each subject. During the focal follows, instantaneous point samples were taken every 2.5 minutes, recording the individual’s state activity and proximity to other monkeys. For example, if the focal individual was engaged in social play at the 2.5 minute mark, “social play” would be recorded for that point sample (although the name of the play partner was not recorded). Only one subject was followed at a time, ensuring that each play instance was only recorded once. Data were collected by one observer who watched and narrated the behaviors, while a second observer input data on a handheld Psion or Android device and assisted observations when necessary (e.g., to confirm the identity of non-focal individuals for the proximity data; Perry et al., 2012). Focal follows were at least 10 minutes in duration, with some longer depending on the year of the study.

While the length of the focal follows varied, the protocol for data collection was otherwise uniform throughout the study period. Consistency between observers was ensured by interobserver reliability tests. Before contributing data, all observers had to pass tests requiring 100% accuracy on monkey identifications, 100% accuracy in the coding scheme, 97% accuracy in speed typing, and 97% accuracy in matching their recorded observations to those of other trained observers in the field during focal follows. Interobserver reliability tests were repeated monthly to ensure lack of drift, and samples for which the observer team (typist and spotter) disagreed regarding monkey identifications or behaviors were discarded.

Observers rotated between the different habituated groups in teams of two or more such that one to three groups were followed simultaneously on a given day, depending on the size of the observation team at the time. The number of days that each group was followed also depended on the size of the observation team, although effort was made to observe each habituated group at least once a month. There was high variability in the number of point samples for each subject per year, ranging from three to 2,456 point samples for years in which data could be collected on each subject (mean = 876.0 samples per monkey/per year, SD = 641.5). This variability is expected to be unbiased across subjects, as it depended primarily on the level of difficulty in locating monkey groups in the wild and varying levels of sampling intensity across the study years.

For the purposes of this study, observers recorded an individual’s behavior as ***social play*** if, at the time of the point sample, the individual was engaged in any of a number of specific play behaviors performed with at least one social partner (e.g., Play Bite, Play Chase, Play Flee, Play Hit, Play Invite/Play Face, Play Bounce/Jump, Play Pull, Play Overlord, Play Push, Play Lunge, Play Threat, Play Pounce On, Play Wrestle, Play Wrestle While Hanging from Tails; see Supporting Information Table 1 for descriptions of behaviors). These behaviors were identified as playful, rather than aggressive, by the presence of play-specific signals (e.g., play-specific facial expressions), absence of audible vocalizations, modified forms (e.g., slow, exaggerated movement, bouncy gait, or gentle versions of aggressive behaviors), or by their co-occurrence with other behaviors that clearly fit those criteria. ***Solitary play*** was recorded if the individual was engaging in object manipulations for no obvious foraging purpose, or engaging in extraneous, sometimes exaggerated body movements that seem to serve no obvious purpose for locomotion, foraging, care of the body (comfort or hygiene), or social interaction. ***Grooming*** was defined as “one monkey picks through the hair of another monkey with the hands and/or mouth; the recipient of this behavior is generally in a reclining posture.”

### Statistical Analysis

To determine sex differences in the rates of each behavior, we ran a series of generalized linear mixed models, with social play, solitary play, and grooming as the outcome variables. Each model included sex, age, and the interaction between sex and age as predictor variables, with a random effect for individual to account for repeated sampling for each individual over time. This random effect allowed the intercepts to vary by individual but not the regression coefficients (i.e., the main effects of sex, age, or sex*age were not allowed to vary by individual).

For each model (social play, solitary play, and grooming), the outcome variable was the count of the total point samples per year in which the given behavior occurred. To achieve this, the point sample data were aggregated for each individual by each year of age. Thus, for each year, an individual had a count of total point samples, and a count of point samples for each behavior (e.g., for the given year, how many point samples that individual was observed in social play, solitary play, and grooming). The total number of point samples per year was included as the exposure variable in the regression models in order to model the outcome variable as a rate and to control for variation in the number of times each individual was observed.

We considered several regression models for each of the three behaviors (social play, solitary play, and grooming) and picked the best models for each behavior based on both AIC and BIC. We report the details of all models considered in the Supporting Information. We first ran all models using a Poisson distribution, which is commonly used to model count variables but which has a strong assumption (that the variance equals the mean). We then compared these original Poisson models to models which estimate the same parameters but assume other negative binomial distributions (Type 1, i.e., nb1 parameterization and Type 2, i.e., nb2 parameterization). The negative binomial models have more relaxed assumptions than Poisson models (by allowing the variance to be greater than the mean). For all three behavioral outcomes, it appeared that the assumptions of the Poisson distribution were violated because the negative binomial distributions provided better model fit (according to AIC and BIC comparisons; see Supporting Information Tables 2, 5, and 8). Type 1 negative binomial provided the best fit for social play, while Type 2 negative binomial provided the best fit for solitary play and grooming.

In order to allow the effect of age on all three behaviors to vary over developmental time, we additionally considered models with a quadratic age predictor variable. Thus, we included both the linear predictor variable, *age*, and a quadratic variable, *age^2^* in each model (social play, solitary play, and grooming), and also included the interactions of both of these age terms with *sex* to allow flexibility in the sex effect over time. Then, the models for each behavior were compared using AIC and BIC to assess whether the addition of the quadratic variable improved model fit. The quadratic age variable improved model fit for grooming and social play, but did not improve model fit for solitary play. Thus, we concluded that the age pattern was better modeled with the quadratic predictor variable only for the social play and grooming models. See the Supporting Information for a list of all the models and their comparison using AIC and BIC.

We were initially concerned that the behavioral outcome variables for our models could be affected by the social group, because different groups could have different numbers of available partners for social interaction, or different play styles. To control for this, we also ran all models with an additional random effect of social group (in this case, the natal group). We found no between-group differences in social play and grooming rates by natal group (variance estimates < 0.0001 in both models), and only a small between-group difference in solitary play rates (variance estimate = 0.129) which did not substantially change the fixed effect estimates on our main variables of interest (sex, age, and the interaction of sex and age) for our best model. This effect may have been partially explained by an association of group size and solitary play rates: larger groups were associated with a slightly lower rate of solitary play, such that for each additional monkey in the group, the rate of solitary play was estimated to be lower by approximately 3 out of 100 point samples per year. Additionally, adding the natal group random effect to the best models for all three behaviors increased AIC and BIC, suggesting that the addition of this parameter did not improve model fit (see SI tables 2, 5, and 8). Because these group effects on the rates of our behaviors of interest were inconsequential and tangential to the main research questions, and model fit was worse when the random effect of natal group was added to the models, we concluded that it was not necessary to add these as control variables to the final models.

All statistical analyses were conducted in the R programming environment (R Core Team, 2019). Regression models were fit using the *glmmTMB* function within the *glmmTMB* package (we originally used *glmer* within the *lme4* package for the Poisson models but found *glmmTMB* was preferrable for the negative binomial models; Bates et al., 2015; Brooks et al., 2017). 95% confidence intervals for the predicted values in all figures were calculated using the *bootMer* function in the *lme4* package. To additionally aid our inferences on the sex differences in behavior at different ages, and because we interacted the sex and age variables, we calculated simple effects for the rate ratios between males and females at different ages and their corresponding p-values (see Supporting Information Tables 4, 7, and 10). Full details of the best models are presented in the Supporting Information (Tables 3, 6, and 9). The data set is provided at https://osf.io/jy3w5/ and the code required to replicate the results are provided at https://osf.io/ybvfg/.

## RESULTS

### Do rates of social play vary by sex and age?

The best social play model used a type 1 negative binomial distribution and included a quadratic age predictor variable. It was given by the following model formula: *socialplay ∼ sex + age_st + sex*age + age^2^ + age^2^*sex + (1 | subject), data=all_data, family=“nbinom1”, offset=log(totalobs)*, where *socialplay* is the count of point samples engaged in social play, *sex* is the subject’s sex (with female as the reference group), *age* is the subject’s age in years (such that zero is the first year of life), *subject* is the unique ID for the subject, *all_data* is the name of our dataset, and *totalobs* is the total count of point samples for the subject-year (all point sample observations for the subject for each year of age). Figure 1 and Supporting Information Table 4 show the model predictions for rates of social play for males (blue) versus females (red) as they change with age. Female rates of social play start out a bit lower than males’ rates and decline steadily with age; males’ rates increase up until their third year of life (when they play at twice the rate of females), and then decline, becoming statistically indistinguishable from female play rates around the tenth year of life, after which the rates for both sexes continue to decline and approach zero. This peak in male play rates roughly corresponds to the juvenile period when males are starting to explore options for co-dispersal from the natal group.

**Figure 1:**
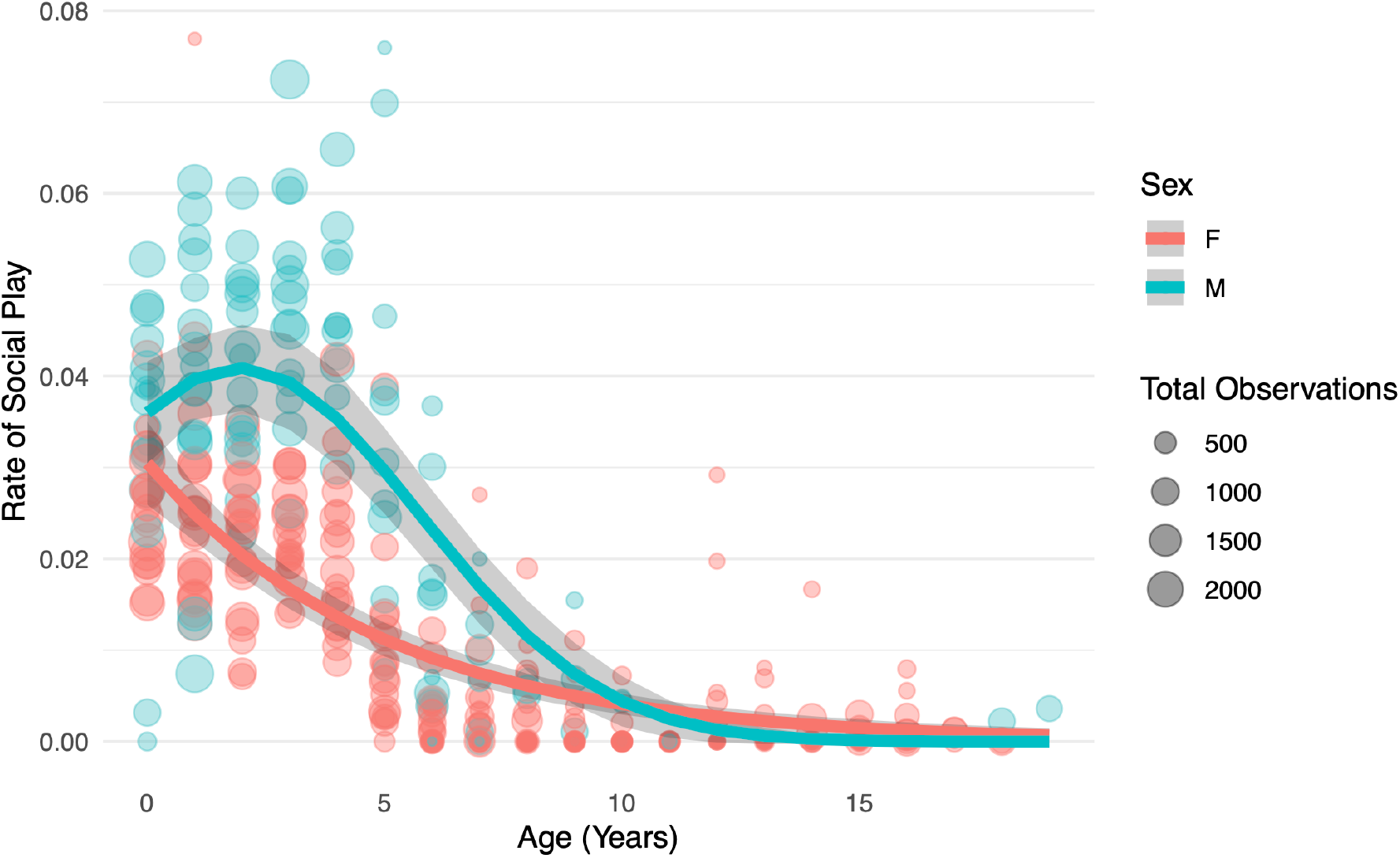
Social Play Model Predictions. Rates of social play (as a proportion of the total point samples per year) are plotted against age in years. The first year of life is coded as 0. Circles represent the proportion of point samples in which an individual monkey was engaging in social play in a given year, with the size of the circle representing the total number of point samples for the individual in the same year. Lines represent predictions from the best regression model chosen from AIC and BIC model comparisons. Shaded areas represent bootstrapped 95% confidence intervals of the predictions.

### Do rates of solitary play vary by sex and age?

The best solitary play model used a type 2 negative binomial distribution and only included the linear age predictor. It was given by the following model formula: *soloplay ∼ sex + age + sex*age + (1 | subject), data=all_data, family=“nbinom2”, offset=log(totalobs).* Figure 2 and Supporting Information Table 7 show the model predictions for solitary play rates for males (blue) versus females (red) as they change with age. Although there was a statistically significant interaction of sex and age, at no age do male and female solitary play rates show biologically meaningful differences from one another. The rate of solitary play declines throughout the juvenile phase, reaching values very close to zero for both sexes by the time they reach adulthood at age 7-10.

**Figure 2:**
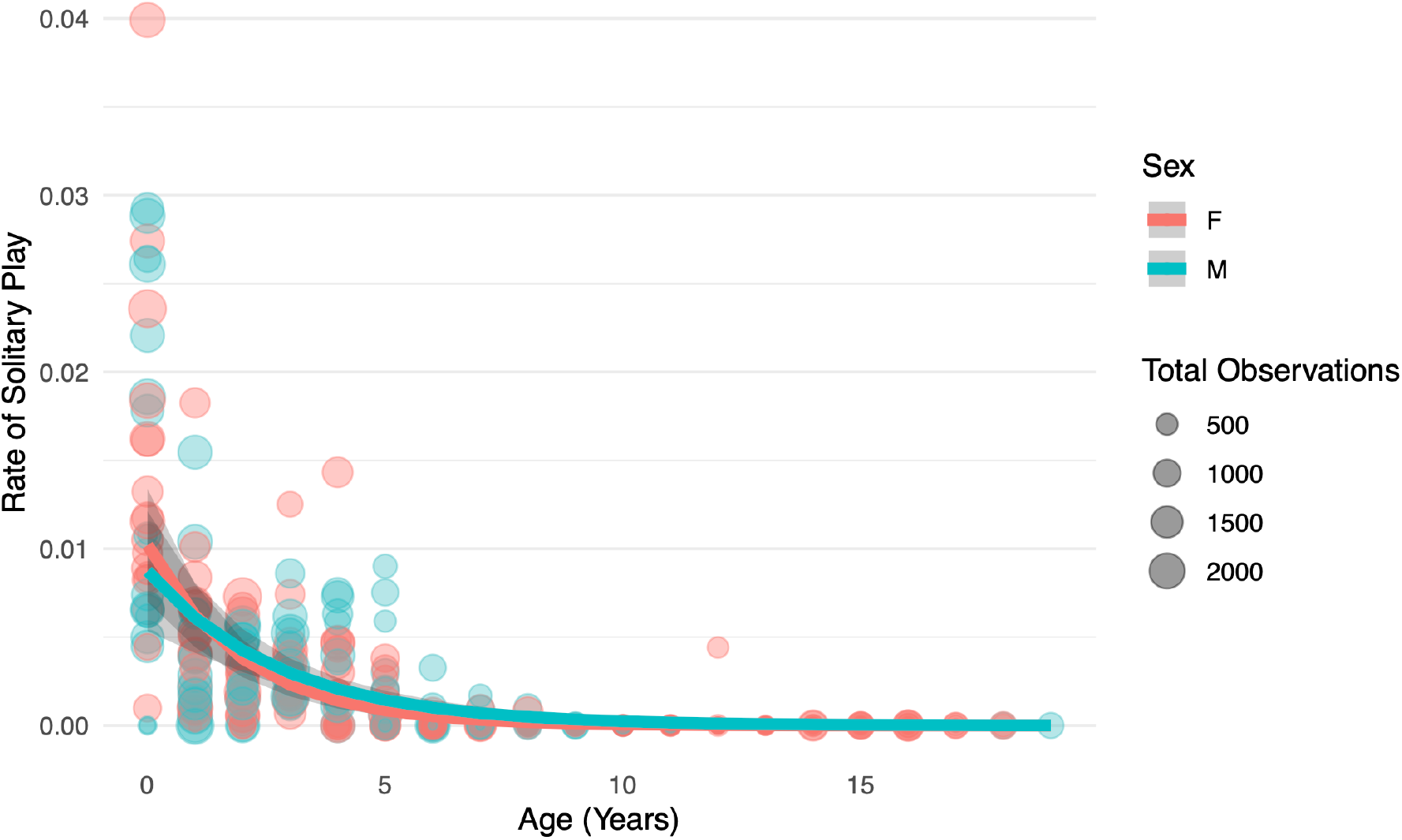
Solitary Play Model Predictions. Rates of solitary play (as a proportion of the total point samples per year) are plotted against age in years. The first year of life is coded as 0. Circles represent the proportion of point samples in which an individual monkey was engaging in solitary play in a given year, with the size of the circle representing the total number of point samples for the individual in the same year. Lines represent predictions from the best regression model chosen from AIC and BIC model comparisons. Shaded areas represent bootstrapped 95% confidence intervals of the predictions. Note that the outlier from Supporting Information Figure 1 is not displayed in this figure.

The solitary play data had one extreme outlier, a six-year-old male (RU) who was observed to have an unusually high rate of solitary play (annual rate of solitary play = 0.2, Z-score among all six-year-olds = 5.1, see Supporting Information Figure 1). However, this extreme value was likely an artifact of the small number of observations for that individual in his sixth year: there were only five point samples for him that year, one of which was solitary play. Because the solitary play regression model accounted for the total number of point samples as the exposure variable, this outlier is not likely to have had an outsized effect on the model overall. In fact, rerunning the model with the outlier removed had no substantive effect on any of the coefficient estimates or inferences (ß_sex_= −0.148, P= 0.532, ß_age_= −0.495, P<0.001, ß_sex*age_= 0.128, P= 0.030). Thus, the outlier was kept in the dataset when fitting the model but removed from Figure 2 for visualization purposes.

### Do rates of grooming vary by sex and age?

The best grooming model used a type 2 negative binomial distribution and included a quadratic age predictor variable. It was given by the following model formula: *grooming ∼ sex + age + sex*age + age^2^ + age^2^*sex + (1 | subject), data=all_data, family=“nbinom2”, offset=log(totalobs)*. Figure 3 and Supporting Information Table 10 show the model predictions for grooming rates (including both giving and receiving grooming) for males (blue) versus females (red) as they change with age. Males are very rarely involved in grooming at any point in their lives. Males start life having grooming rates that are indistinguishable from those of females. By the third year of life, females already show higher rates of grooming than males, and they continue to increase the proportion of time they spend grooming until around the eleventh year of life, at which point their grooming rates start declining again, although it remains higher than the male rate.

**Figure 3:**
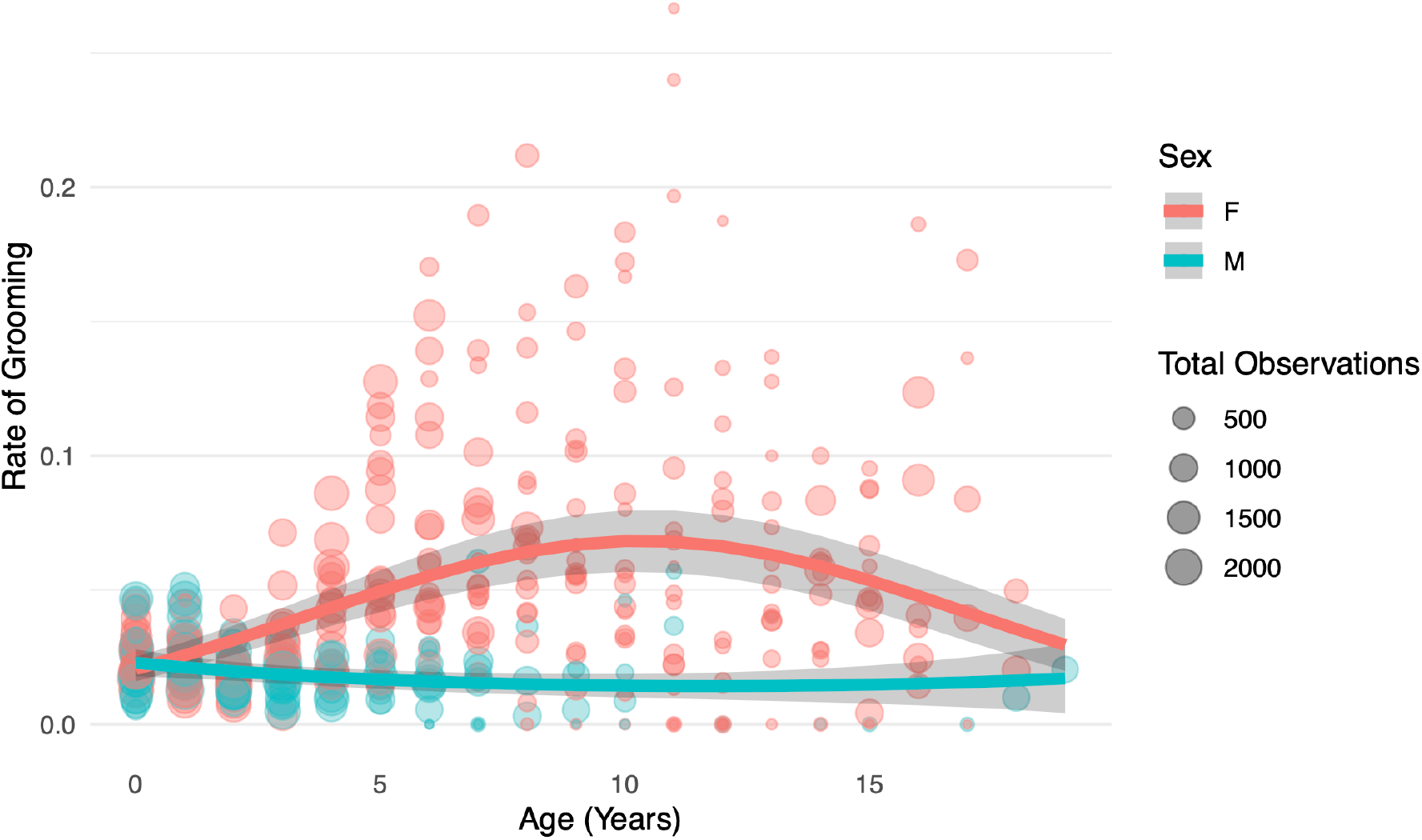
Grooming Model Predictions. Rates of grooming (as a proportion of the total point samples per year) are plotted against age in years. The first year of life is coded as 0. Circles represent the proportion of point samples in which an individual monkey was engaging in grooming in a given year, with the size of the circle representing the total number of point samples for the individual in the same year. Lines represent predictions from the best regression model chosen from AIC and BIC model comparisons. Shaded areas represent bootstrapped 95% confidence intervals of the predictions.

## DISCUSSION

Consistent with the general mammalian trend, rates of play in white-faced capuchins were greatest early in life and declined with age, for both social and solitary play and for both males and females. There was a sex difference in the rates of social play in this species, such that the average rate was higher for juvenile males than females, although the sex difference disappears in adulthood. Additionally, while females had declining rates of social play as age increased, rates of social play in males actually increased from ages zero to two, before declining with age. However, our analysis of solitary play found that the predicted rates of solitary play did not differ meaningfully between males and females. We found only a negligible (albeit statistically significant) interaction effect between sex and age, such that females had a slightly steeper decline of solitary play with age, compared to males. Finally, we analyzed trends in grooming behavior to investigate the possibility of time budget tradeoffs between social play and grooming, a known bond-formation behavior. Male capuchins participated in grooming at very low rates throughout life, while females groomed at higher rates, with a peak in grooming rates around age eleven. These patterns of grooming were consistent with previous findings on grooming in white-faced capuchins which suggest that adult females participate in grooming much more often than adult males (Manson et al., 1999; Perry, 1996, 1998).

Together, these results suggest that white-faced capuchin males allocate more time toward social play than females, especially during the juvenile period. Interestingly, the interaction of sex and age on social play rate appears in Figure 1 to be at least partially driven by males (but not females) increasing rates of social play over time during the early juvenile period, with male social play expected to peak around ages 1-3. Therefore, males maximize social play at a time just before the average age of dispersal out of the natal group. This suggests that social play may have particular benefits for male survival or reproductive success during the risky dispersal period.

Under the practice hypothesis, playing provides benefits for the development of sex-specific behaviors. The finding of higher rates of social play, but not solitary play, in juvenile male white-faced capuchins, compared to females, could reflect important differences in the types of social behaviors that are most crucial for each sex’s reproductive success in adulthood. Fighting skills are important for male reproductive success as they allow males to enter and take over groups of females. Males that are unable to achieve alpha status in new social groups might never gain access to reproductive opportunities. Fighting requires not just physical agility, as might be developed through solitary play, but also tactical maneuvers and social skills, which likely require practice with partners. Social play may also help to develop a greater understanding of the social affordances of physical maneuvers—for example, assessing one’s own physical power relative to others, predicting the loyalty of allies, or learning how to avoid conflict escalation. In contrast, physical fighting skills are not as critical for female reproductive success. If social play allows white-faced capuchins to explore and hone specific skills that are needed in adulthood, our finding that males partake in higher rates of social (primarily rough-and tumble) play are consistent with the practice hypothesis.

The results are also consistent with the bonding hypothesis for the function of play, which is not necessarily mutually exclusive from the practice hypothesis. We propose that females may compensate for lower opportunities to form social bonds via play through increases in their rates of grooming. This sex difference in development may simply reflect that females and males allocate social effort in different ways, or it may reflect that males and females have different needs for developing the particular kinds of tactical skills that can be practiced in rough-and-tumble social play. If social play and grooming both function to form social bonds, our results could be consistent with females preferentially forming social bonds through grooming rather than play, compared to males. However, because we did not directly analyze the relationship between social play and grooming, more research would be needed to substantiate this exploratory association. Additionally, future research might explore the role of socioecological factors in explaining why grooming rates are highest in adulthood for females, rather than during the juvenile period as is the case for male social play.

This interpretation of our results with respect to the bonding hypothesis assumes that both sexes are preferentially grooming and playing with same-sex partners. However, the point samples analyzed for this study did not contain information on the partners or direction of social behaviors. For example, whether grooming was given or received at the time of the point sample was not specified, so we are left to conjecture the probable biases in directions based on previous studies of patterning of grooming in adults. Previous studies have shown that adult female white-faced capuchins are most likely to groom with other females or the alpha male (Perry 1996). Adult males at Lomas Barbudal rarely groom one another, and when they do participate in grooming interactions, it is often when they receive grooming from females (Perry 1997, 1998). In fact, it is likely that a large portion of the grooming time attributed to adult males in this study was due to males (particularly alpha males) receiving grooming from females. Previous research also suggests that preferences for social play partners may depend on sex in some primate species (e.g., Lutz et al., 2019; Maestripieri & Ross, 2004).

An additional limitation of our dataset for this study is that although there was a distinction between social and solitary play, there were no distinctions between subtypes of these categories. For example, the social play category included all rough-and-tumble play behaviors, and distinctions were not made between specific behaviors within that category (e.g., play bouncing versus play biting; see ethogram in Supporting Information Table 1). The solitary play category included both object and locomotor play. An additional challenge with recording solitary, object play was that a conservative definition was used during data collection that excluded handling of leaves, sticks, or other food items in the absence of any obvious repetition, modification, or exaggeration from functional food handling. Thus, handling of food items was generally categorized as foraging in the dataset, even though theoretically a monkey could play with those items. Most definitions of play in the animal literature exclude behaviors that could be interpreted as foraging, so the exclusion of these behaviors is not unique to the current study (Burghardt, 2005). However, it is important to recognize that studies of play in wild animals probably underestimate the rate of object play by disregarding play that occurs with food items, particularly for omnivorous animals like capuchins (in which case even inedible objects like sticks or rocks are commonly handled to eat insects off of them).

This study raises several questions that should be explored in future work using continuous, rather than instantaneous, sampling of play behavior. Similarly to how sex-specific partner preferences may influence rates of social play and grooming, the rate of social play might also depend on the number of siblings or the total number of juveniles in each group that are available as playmates at any given time. The relationship of play to social bonding should also be investigated further. For example, if male white-faced capuchins that play together as juveniles are more likely to co-disperse together, that could indicate that play provides important opportunities for young males to strengthen and test these fitness-relevant relationships, while in the protected space of their natal group. While Perry and colleagues (2017) found no relationship between rates of social play and time to emigration or time to first alpha male status, that paper used a broader sample of individuals with lower sampling rates, and did not consider the identity of play or co-emigration partners. The bonding hypothesis might also be supported if females that play together as juveniles are more likely to support each other in aggressive coalitions, or to be grooming partners in adulthood. Future comparisons between species may also give us clues into the bonding functions of play. For example, social bonds may be particularly important for capuchins, resulting in both physical and bonding benefits of play, whereas in species where there is less reproductive skew or where reproductive success is less dependent on bonds and alliances, play (even social play) may primarily confer physical benefits. Finally, longitudinal datasets such as the one used in this study provide excellent opportunities to investigate whether play in early life is associated with any concrete fitness benefits in adulthood. With longitudinal data, it may be possible to assess delayed fitness benefits in variables like rank acquisition, number of offspring, or outcomes linked to fighting skills such as fighting success or wounding frequency.

While the current study findings are consistent with several functional hypotheses regarding play, it is important to remember that sex differences in play are not *necessarily* meaningful in terms of adult behavior. For example, a study in meerkats found no correlation between the frequency of play-fighting in early life and subsequent fighting success in adulthood, for either sex (Sharpe, 2005). Cords and colleagues (2010) have argued that sex differences in juvenile behavior can anticipate adult behavior in some cases, but researchers should not overlook the immediate consequences of a behavior in favor of delayed ones. Additionally, Pellegrini and Smith (2005) argued that although it is possible that males and females in many species evolved different play styles to develop different skills needed in adulthood, it is also possible that sex differences in juvenile play could simply be due to size dimorphisms or to sex-specific nutritional or energetic demands at different points in development.

This longitudinal study of the development of play behavior in wild capuchins provides support for both the practice and bonding hypotheses for the evolution of play behavior. These results demonstrate that males engage in more social play than females do, especially as juveniles, but that there is essentially no sex difference in the rates of solitary play (which is hypothesized to prepare individuals for foraging and locomotor skills equally critical to both sexes). Rates of both social and solitary play generally decrease as capuchins age into adulthood. The trajectories of age-related change in social play and grooming can perhaps tell us something about the development of sex-specific social strategies. Males’ rates of social play decline less rapidly than females’ and even show a slight increase shortly before the age at which males typically disperse to new social groups. Females’ grooming rates increase over the course of early development, whereas males’ grooming rates decrease. These patterns seem to reflect preparation for distinct male and female strategies for increasing reproductive success, in which females use grooming as a way to service their social relationships, and males use rough-and-tumble play to practice fighting and forge alliances with other males, whose coalitionary aid is essential for them to obtain breeding positions.

## ACKNOWLEDGEMENTS

Thank you to A. Lin, D. Cohen, K. Kajokaite, and S. Jalal for their invaluable assistance with the statistical analyses. Thanks also to E. Cartmill, H. C. Barrett, and M. Cooper-White for helpful comments on the manuscript drafts. The following field assistants assisted S. Perry in data collection by contributing a year or more of data to the Lomas Barbudal Monkey Project dataset: C. Angyal, A. Autor, B. Barrett, L. Beaudrot, M. Bergstrom, R. Berl, A. Bjorkman, L. Blankenship, T. Borcuch, J. Broesch, D. Bush, J. Butler, F. Campos, C. Carlson, M. Corrales, J. Damm, C. de Rango, C. Dillis, N. Donati, G. Dower, R. Dower, A. Duchesneau, K. Feilen, S. Fiello, K. Fisher, A. Fuentes J., M. Fuentes A., C. Gault, H. Gilkenson, I. Godoy, I. Gottlieb, J. Griciute, J. Gros-Louis, L.M. Guevara, L. Hack, R. Hammond, S. Herbert, C. Hirsch, M. Hoffman, A. Hofner, C. Holman, S. Hyde, O. Jacobson, W. Lammers, L. Johnson, S. Lee, S. Leinwand, S. Lopez, T. Lord, S. MacCarter, M. Kay, E. Kennedy, D. Kerhoas-Essens, E. Johnson, S. Kessler, J. Mackenzie, J. Manson, F. McKibben, W. Meno, A. Mensing, C. Mitchell, Y. Namba, A. Neyer, C. O’Connell, J.C.Ordoñez J., N. Parker, B. Pav, R. Popa, K. Potter, K. Ratliff, N. Roberts Buceta, H. Ruffler, S. Sanford, M. Saul, I. Schamberg, N. Schleissmann, C. Schmitt, A. Scott, J. Shih, K. van Atta, L. van Zuidam, J. Verge, A. Walker-Bolton, E. Wikberg, E. Williams, L.E. Wolf, and D. Wood. Particular thanks is owed to H. Gilkenson and W. Lammers for long-term management of the site during 2002-2013. D. Cohen assisted in development of the database and writing of queries.

Funding for this project was provided by the UCLA Graduate Summer Research Mentorship program and the UCLA Anthropology department. Funding for data collection was provided by MPI-EVAN, UCLA and several grants to S. Perry: NSF (BCS-1919649, BCS-1638428, BCS-0613226, BCS-848360), National Geographic Society (7968-06, 8671-09, 20113909, 9795-15, 45176R-18), Templeton World Charity Foundation (0208), and five Leakey Foundation grants. S. Winkler is supported by the NSF Graduate Research Fellowship Program (Grant DGE-1650604 and DGE-2034835). Research permission was granted to S. Perry by SINAC and MINAE (the Costa Rican Park Service) and the landowners of Brin d’Amor, Hacienda Pelon de la Bajura, and San Ramon de Bagaces.

## Supporting information

### Information about social groups

All subjects were born into groups AA, FF, RR, and FL. Over the course of the study, AA fissioned into AA, FL (in 2003), and CE (in 2012). FF fissioned into FF and RF (in 2007). RR fissioned into RR, MK (in 2004), DI (in 2012), SP in (1999-2000), and LB (in 2010). MK later fissioned into MK and CU (in 2007). The original name was kept by the larger group after each fission event.

**Supporting Information Table 1:**
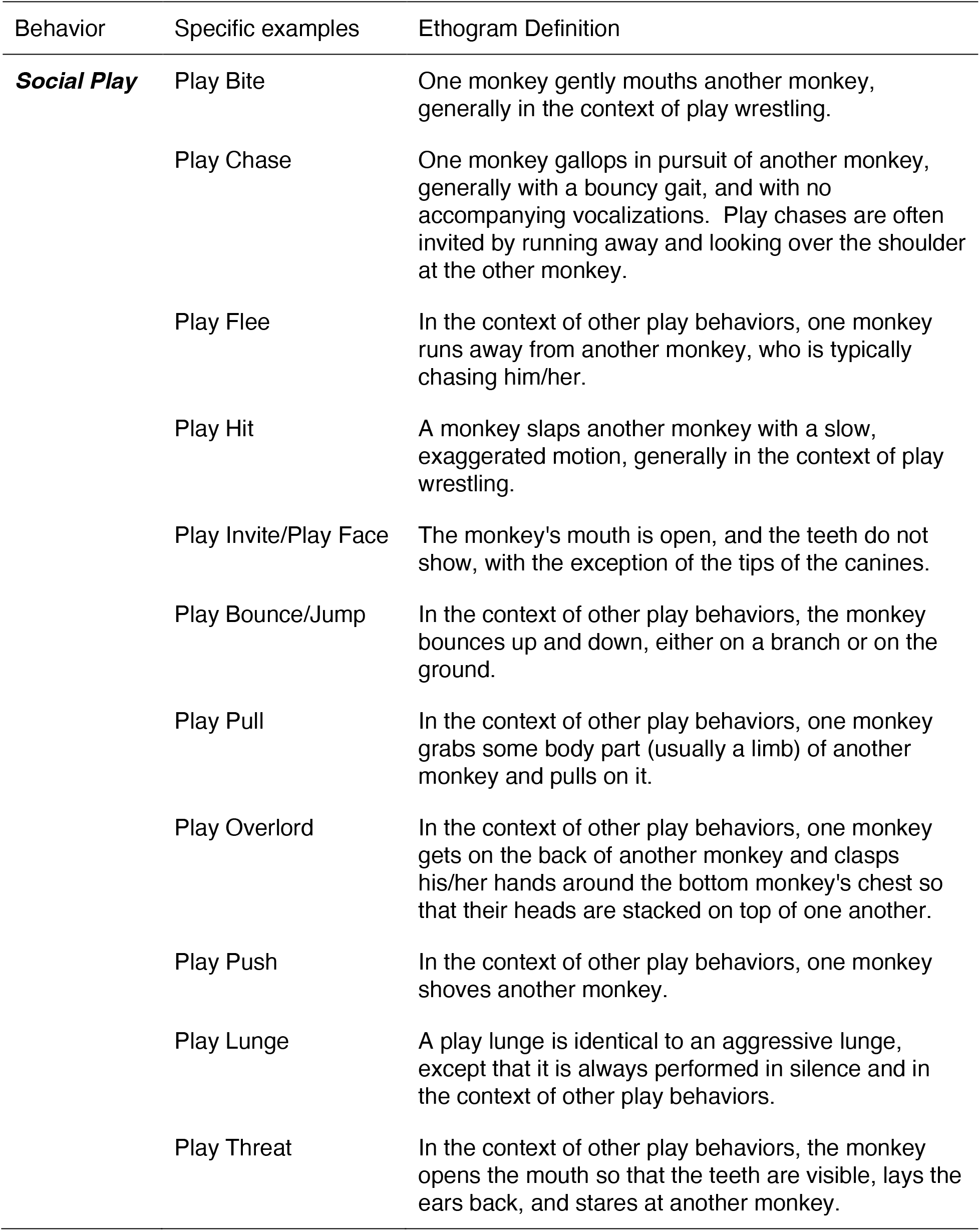
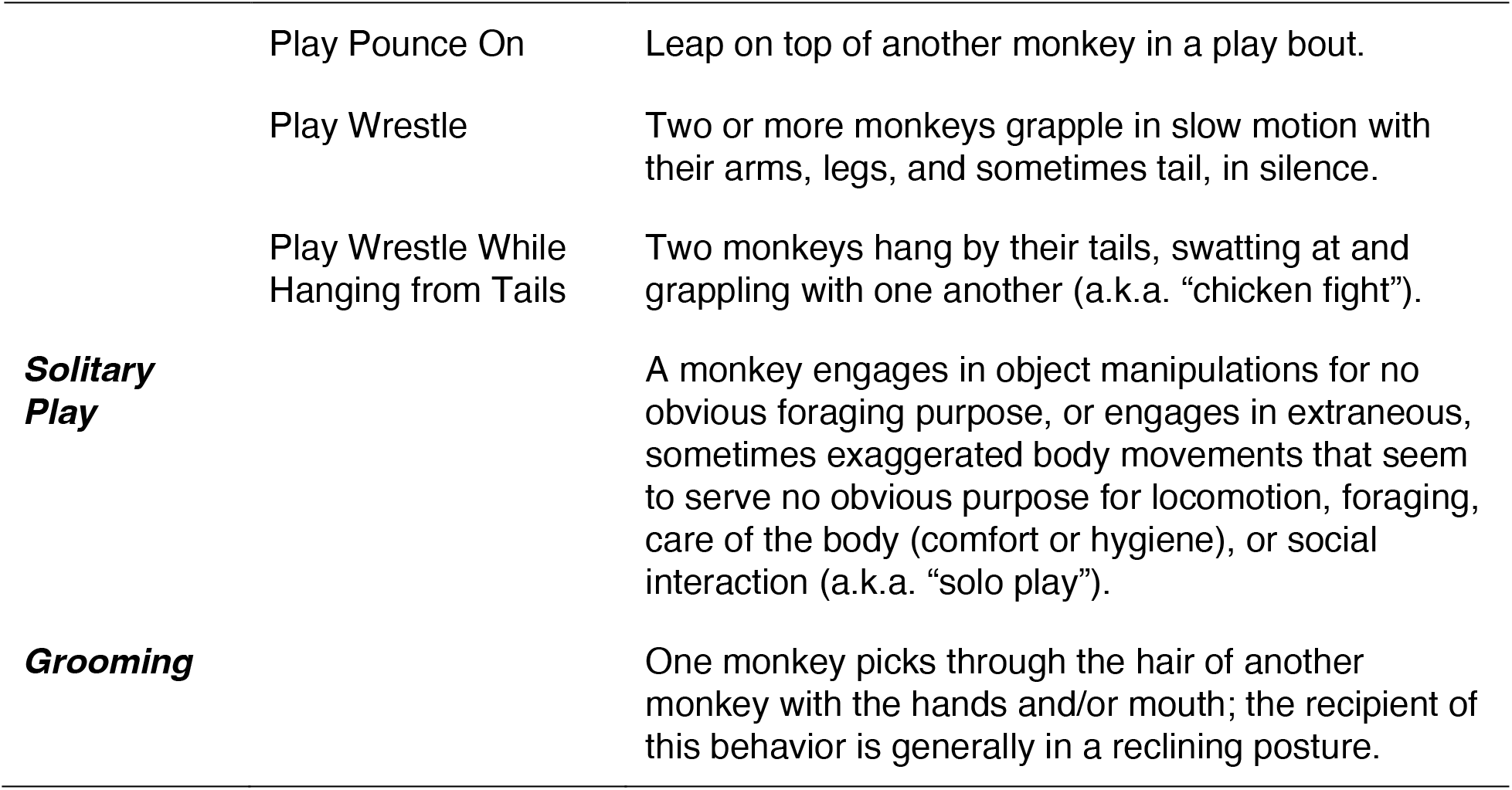
Ethogram of Behaviors.

**Supporting Information Table 2:** All Models for Social Play, with Best Model in Bold.

| Model object name | Model formula | df | AIC | BIC |
| --- | --- | --- | --- | --- |
| mod_sp_poisson | glmmTMB(socialplay ~ sex + age + sex*age + (1 \| subject),<br>data=all_data,<br>family="poisson",<br>offset=log(totalobs)) | 5 | 3164.439 | 3184.434 |
| mod_sp_nbinom1 | glmmTMB(socialplay ~ sex + age + sex*age + (1 \| subject),<br>data=all_data,<br>family="nbinom1",<br>offset=log(totalobs)) | 6 | 2213.815 | 2237.809 |
| mod_sp_nbinom2 | glmmTMB(socialplay ~ sex + age + sex*age + (1 \| subject),<br>data=all_data,<br>family="nbinom2",<br>offset=log(totalobs)) | 6 | 2253.180 | 2277.173 |
| mod_sp_quad_poisson | glmmTMB(socialplay ~ sex + age + sex*age + age2 + age2*sex + (1 \| subject),<br>data=all_data,<br>family="poisson",<br>offset=log(totalobs)) | 7 | 2822.432 | 2850.425 |
| **mod_sp_quadandnbinom1** | **glmmTMB(socialplay ~ sex + age + sex\*age + age2 + age2\*sex + (1 \| subject),**<br>**data=all_data,**<br>**family="nbinom1",**<br>**offset=log(totalobs))** | **8** | **2190.446** | **2222.437** |
| mod_sp_quadandnbinom2 | glmmTMB(socialplay ~ sex + age + sex*age + age2 + age2*sex + (1 \| subject),<br>data=all_data,<br>family="nbinom2",<br>offset=log(totalobs)) | 8 | 2249.053 | 2281.045 |
| mod_sp_quadandnbinom1_group | glmmTMB(socialplay ~ sex + age + sex*age + age2 + age2*sex + (1 \| Natal_group / subject),<br>data=all_data,<br>family="nbinom1",<br>offset=log(totalobs)) | 9 | 2192.446 | 2228.436 |

**Supporting Information Table 3:** Social Play Best Model Results.

| *Dependent variable:*<br>Social Play Observations |  |  |  |  |
| --- | --- | --- | --- | --- |
|  | Estimate | Std. Error | Z Value | P Value |
| Sex (Male) | 0.1663 | 0.1076 | 1.54 | 0.122 |
| Age (Years) | -0.2011 | 0.03205 | -6.27 | 3.54e-10 |
| Age<sup>2</sup> | 9.968e-6 | 0.00278 | 0.00 | 0.9972 |
| Sex*Age | 0.3335 | 0.06033 | 5.53 | 3.26e-8 |
| Sex*Age<sup>2</sup> | -0.03426 | 0.007569 | -4.53 | 6.01e-6 |
| Constant | -3.491 | 0.07352 | -47.48 | <2e-16 |
| Random effect<br>(Subject) | Variance<br>0.0238 | Std. Dev<br>0.1543 |  |  |
| Observations (# subject-<br>years) | 403 |  |  |  |
| Groups (# subject animals) | 38 |  |  |  |
| Log Likelihood | -1087.2 |  |  |  |
| Akaike Inf. Crit. | 2190.4 |  |  |  |
| Bayesian Inf. Crit | 2222.4 |  |  |  |

**Model formula:** mod\_sp\_quadandnbinom1 = glmmTMB(socialplay ~ sex + age + sex\*age + age2 + age2\*sex + (1 | subject), data=all\_data, family="nbinom1", offset=log(totalobs))
**Dispersion parameter for nbinom1 family:** 5.27

**Supporting Information Table 4:** Predicted Values and Simple Effects from Best Social Play Model. The maximum predicted values for each column are in bold.

| Age | Predicted<br>rate for<br>females | Predicted<br>rate for<br>males | Sex difference<br>in rate<br>prediction (M-F) | Rate ratio<br>(M/F) | P-value for<br>the rate ratio |
| --- | --- | --- | --- | --- | --- |
| 0 | **0.0305** | 0.036 | 0.00551 | 1.18 | 0.123 |
| 1 | 0.0249 | 0.0397 | 0.0148 | 1.59 | 1.44E-08 |
| 2 | 0.0204 | **0.0409** | 0.0205 | 2.01 | 8.45E-18 |
| 3 | 0.0167 | 0.0393 | **0.0227** | 2.36 | 1.68E-21 |
| 4 | 0.0136 | 0.0353 | 0.0217 | 2.59 | 9.39E-22 |
| 5 | 0.0112 | 0.0296 | 0.0185 | 2.66 | 1.24E-19 |
| 6 | 0.00912 | 0.0232 | 0.0141 | 2.54 | 1.04E-14 |
| 7 | 0.00746 | 0.017 | 0.00951 | 2.27 | 1.81E-08 |
| 8 | 0.0061 | 0.0116 | 0.00549 | 1.9 | 0.000697 |
| 9 | 0.00499 | 0.00739 | 0.0024 | 1.48 | 0.119 |
| 10 | 0.00408 | 0.0044 | 0.000319 | 1.08 | 0.821 |
| 11 | 0.00334 | 0.00245 | -0.000892 | 0.733 | 0.471 |
| 12 | 0.00273 | 0.00127 | -0.00146 | 0.465 | 0.161 |
| 13 | 0.00224 | 0.000617 | -0.00162 | 0.276 | 0.0573 |
| 14 | 0.00183 | 0.000279 | -0.00155 | 0.153 | 0.0227 |
| 15 | 0.0015 | 0.000118 | -0.00138 | 0.0789 | 0.0101 |
| 16 | 0.00122 | 0.0000466 | -0.00118 | 0.0381 | 0.00507 |
| 17 | 0.001 | 0.0000172 | -0.000984 | 0.0172 | 0.0028 |
| 18 | 0.000819 | 0.00000592 | -0.000813 | 0.00722 | 0.00168 |
| 19 | 0.00067 | 0.0000019 | -0.000668 | 0.00284 | 0.00108 |

**Supporting Information Table 5:** All Models for Solitary Play, with Best Model in Bold.

| Model object name | Model formula | df | AIC | BIC |
| --- | --- | --- | --- | --- |
| mod_wp_poisson | glmmTMB(solo play ~ sex + age + sex*age + (1 \| subject), data=all_data, family="poisson", offset=log(totalobs)) | 5 | 1321.402 | 1341.396 |
| mod_wp_nbinom1 | glmmTMB(solo play ~ sex + age + sex*age + (1 \| subject), data=all_data, family="nbinom1", offset=log(totalobs)) | 6 | 1167.629 | 1191.622 |
| **mod_wp_nbinom2** | **glmmTMB(solo play ~ sex + age + sex\*age + (1 \| subject), data=all_data, family="nbinom2", offset=log(totalobs))** | **6** | **1152.304** | **1176.298** |
| mod_wp_quad_poisson | glmmTMB(solo play ~ sex + age + sex*age + age2 + age2*sex + (1 \| subject), data=all_data, family="poisson", offset=log(totalobs)) | 7 | 1316.548 | 1344.540 |
| mod_wp_quadandnbinom1 | glmmTMB(solo play ~ sex + age + sex*age + age2 + age2*sex + (1 \| subject), data=all_data, family="nbinom1", offset=log(totalobs)) | 8 | 1169.274 | 1201.265 |
| mod_wp_quadandnbinom2 | glmmTMB(solo play ~ sex + age + sex*age + age2 + age2*sex + (1 \| subject), data=all_data, family="nbinom2", offset=log(totalobs)) | 8 | 1156.274 | 1188.265 |
| mod_wp_nbinom2_group | glmmTMB(solo play ~ sex + age + sex*age + (1 \| Natal_group / subject), data=all_data, family="nbinom2", offset=log(totalobs)) | 7 | 1153.217 | 1181.210 |

**Supporting Information Table 6:**
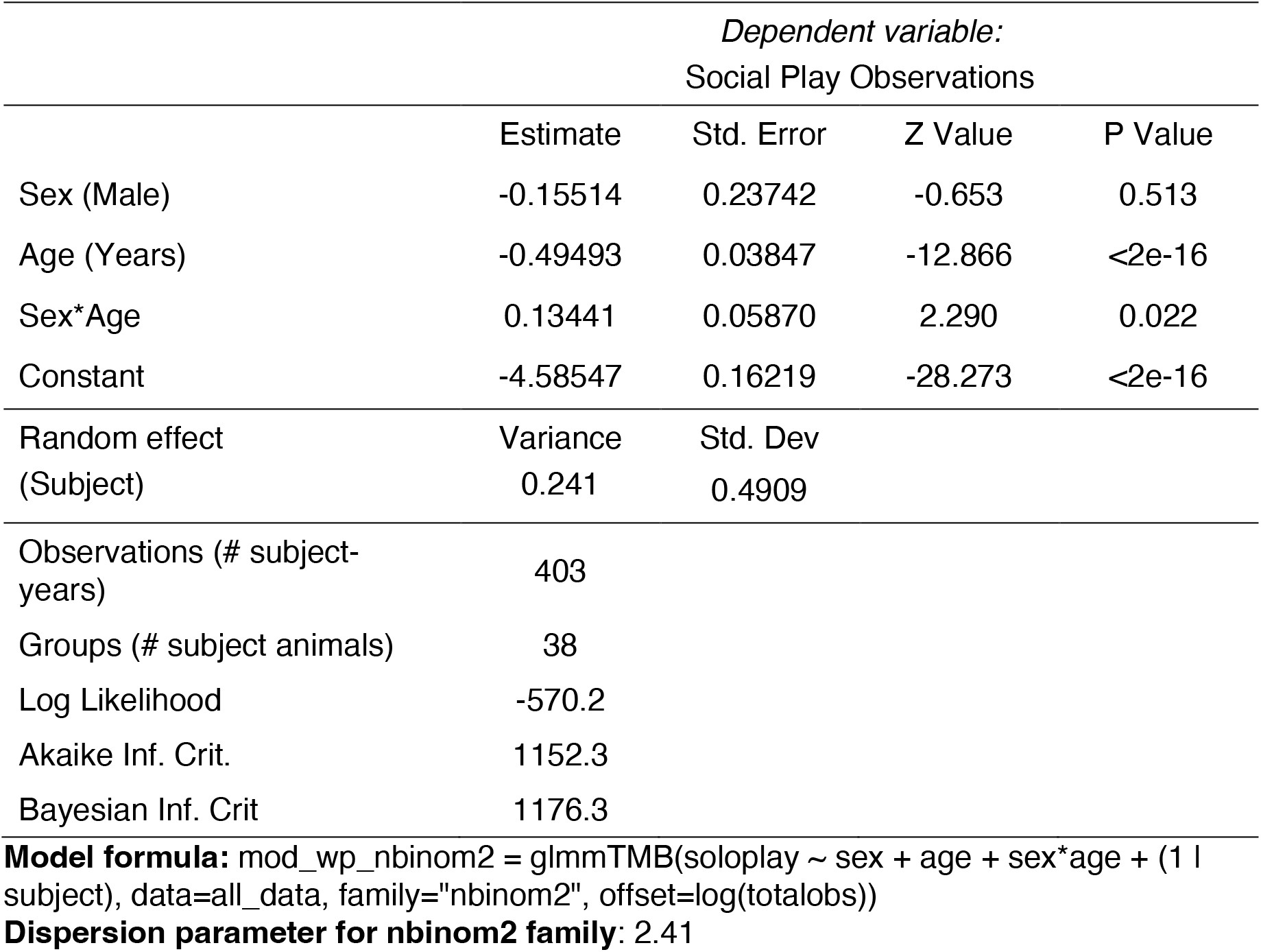
Solitary Play Best Model Results.

**Supporting Information Table 7:** Predicted Values and Simple Effects from Best Solitary Play Model. The maximum predicted values for each column are in bold.

| Age | Predicted rate for females | Predicted rate for males | Sex difference in rate prediction (M-F) | Rate ratio (M/F) | P-value for the rate ratio |
| --- | --- | --- | --- | --- | --- |
| 0 | **0.0102** | **0.00873** | **-0.00147** | 0.856 | 0.513 |
| 1 | 0.00622 | 0.00609 | -0.000128 | 0.979 | 0.922 |
| 2 | 0.00379 | 0.00425 | 0.000456 | 1.12 | 0.573 |
| 3 | 0.00231 | 0.00296 | 0.000651 | 1.28 | 0.232 |
| 4 | 0.00141 | 0.00206 | 0.000656 | 1.47 | 0.0948 |
| 5 | 0.000859 | 0.00144 | 0.000581 | 1.68 | 0.0485 |
| 6 | 0.000523 | 0.001 | 0.000481 | 1.92 | 0.0316 |
| 7 | 0.000319 | 0.0007 | 0.000381 | 2.19 | 0.0244 |
| 8 | 0.000195 | 0.000488 | 0.000294 | 2.51 | 0.0209 |
| 9 | 0.000119 | 0.00034 | 0.000222 | 2.87 | 0.0191 |
| 10 | 0.0000723 | 0.000237 | 0.000165 | 3.28 | 0.0181 |
| 11 | 0.0000441 | 0.000166 | 0.000121 | 3.76 | 0.0176 |
| 12 | 0.0000269 | 0.000115 | 0.0000886 | 4.3 | 0.0173 |
| 13 | 0.0000164 | 0.0000805 | 0.0000641 | 4.91 | 0.0172 |
| 14 | 0.00000998 | 0.0000561 | 0.0000461 | 5.62 | 0.0172 |
| 15 | 0.00000609 | 0.0000391 | 0.0000331 | 6.43 | 0.0172 |
| 16 | 0.00000371 | 0.0000273 | 0.0000236 | 7.36 | 0.0172 |
| 17 | 0.00000226 | 0.000019 | 0.0000168 | 8.41 | 0.0173 |
| 18 | 0.00000138 | 0.0000133 | 0.0000119 | 9.62 | 0.0174 |
| 19 | 0.00000084 | 0.00000925 | 0.00000841 | 11 | 0.0175 |

**Supporting Information Figure 1:**
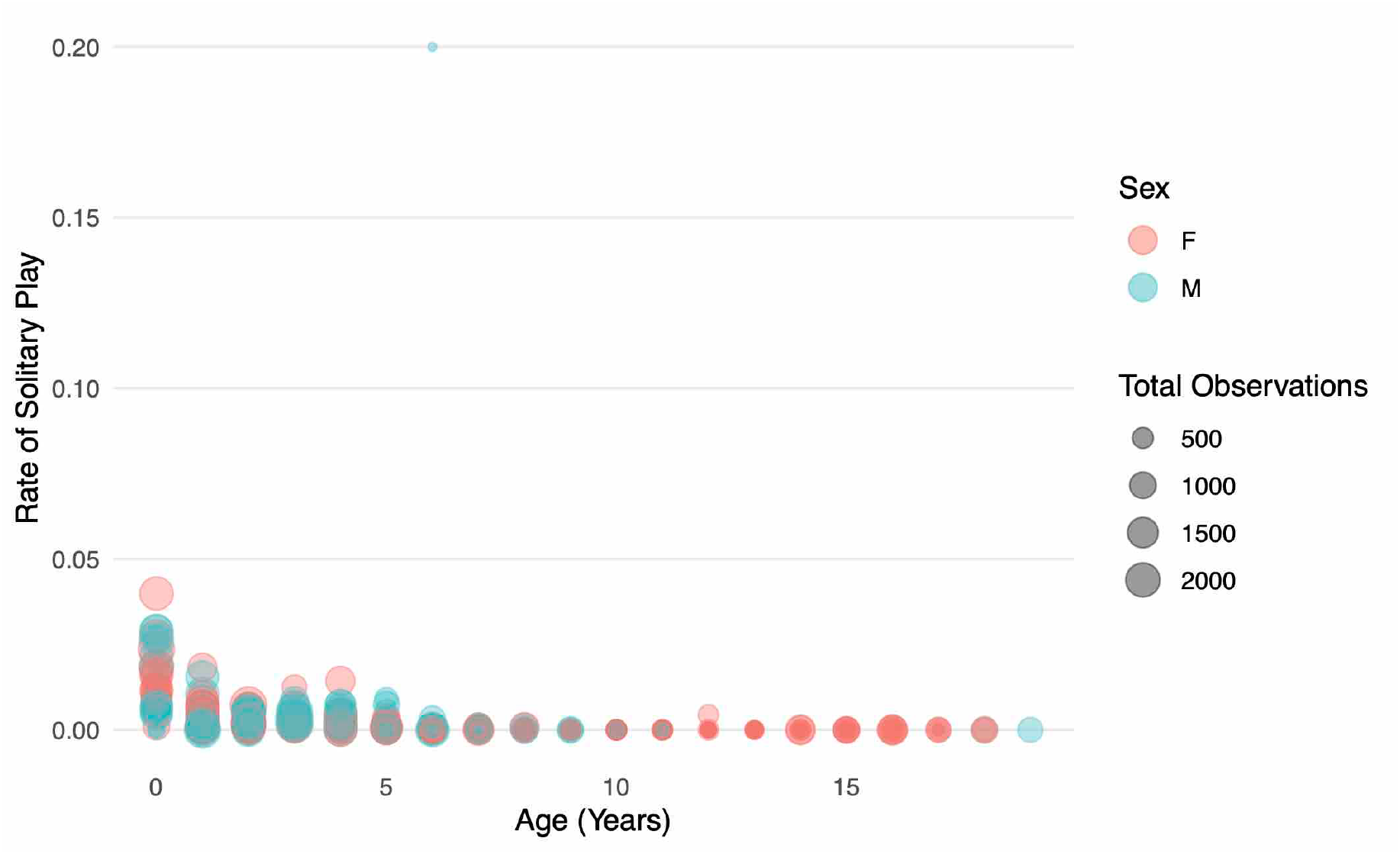
Solitary Play Rates with Outlier. Rates of solitary play (as a proportion of the total point samples per year) are plotted against age in years. The first year of life is coded as 0. Circles represent the proportion of point samples in which an individual monkey was engaging in solitary play in a given year, with the size of the circle representing the total number of point samples for the individual in the same year. In this figure, the outlier (male with solitary play in 1 of 5 point samples taken at age 6) can be clearly seen. The outlier was included in all analyses but removed from Figure 2 in order to better visualize the model predictions.

**Supporting Information Table 8:**
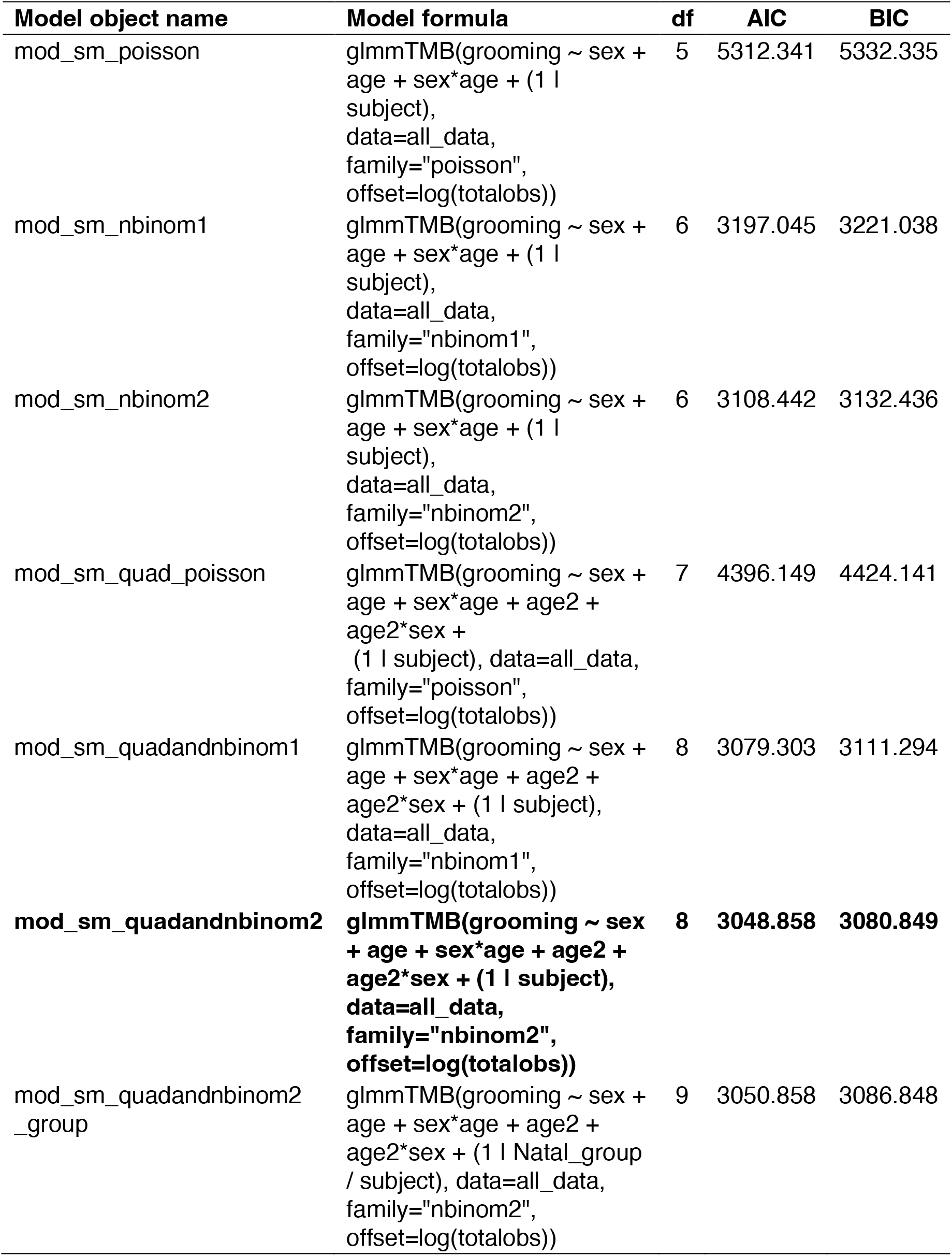
All Models for Grooming, with Best Model in Bold.

| Model object name | Model formula | df | AIC | BIC |
| --- | --- | --- | --- | --- |
| mod_sm_poisson | glmmTMB(grooming ~ sex + age + sex*age + (1 \| subject), data=all_data, family="poisson", offset=log(totalobs)) | 5 | 5312.341 | 5332.335 |
| mod_sm_nbinom1 | glmmTMB(grooming ~ sex + age + sex*age + (1 \| subject), data=all_data, family="nbinom1", offset=log(totalobs)) | 6 | 3197.045 | 3221.038 |
| mod_sm_nbinom2 | glmmTMB(grooming ~ sex + age + sex*age + (1 \| subject), data=all_data, family="nbinom2", offset=log(totalobs)) | 6 | 3108.442 | 3132.436 |
| mod_sm_quad_poisson | glmmTMB(grooming ~ sex + age + sex*age + age2 + age2*sex + (1 \| subject), data=all_data, family="poisson", offset=log(totalobs)) | 7 | 4396.149 | 4424.141 |
| mod_sm_quadandnbinom1 | glmmTMB(grooming ~ sex + age + sex*age + age2 + age2*sex + (1 \| subject), data=all_data, family="nbinom1", offset=log(totalobs)) | 8 | 3079.303 | 3111.294 |
| **mod_sm_quadandnbinom2** | **glmmTMB(grooming ~ sex + age + sex\*age + age2 + age2\*sex + (1 \| subject), data=all_data, family="nbinom2", offset=log(totalobs))** | **8** | **3048.858** | **3080.849** |
| mod_sm_quadandnbinom2_group | glmmTMB(grooming ~ sex + age + sex*age + age2 + age2*sex + (1 \| Natal_group / subject), data=all_data, family="nbinom2", offset=log(totalobs)) | 9 | 3050.858 | 3086.848 |

**Supporting Information Table 9:**
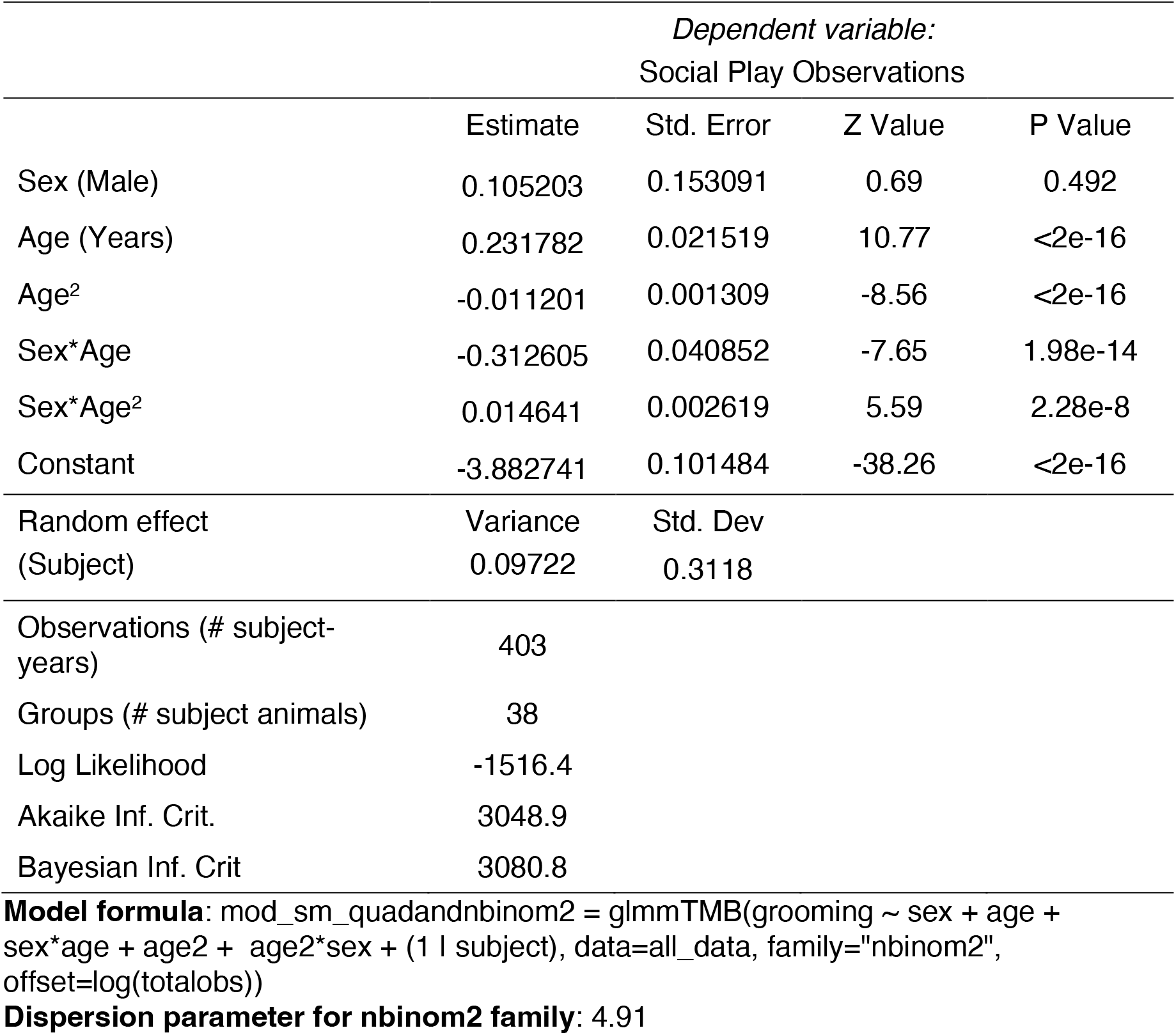
Grooming Best Model Results.

| *Dependent variable:*<br>Social Play Observations |  |  |  |  |
| --- | --- | --- | --- | --- |
|  | Estimate | Std. Error | Z Value | P Value |
| Sex (Male) | 0.105203 | 0.153091 | 0.69 | 0.492 |
| Age (Years) | 0.231782 | 0.021519 | 10.77 | <2e-16 |
| Age<sup>2</sup> | -0.011201 | 0.001309 | -8.56 | <2e-16 |
| Sex*Age | -0.312605 | 0.040852 | -7.65 | 1.98e-14 |
| Sex*Age<sup>2</sup> | 0.014641 | 0.002619 | 5.59 | 2.28e-8 |
| Constant | -3.882741 | 0.101484 | -38.26 | <2e-16 |
| Random effect<br>(Subject) | Variance<br>0.09722 | Std. Dev<br>0.3118 |  |  |
| Observations (# subject-<br>years) | 403 |  |  |  |
| Groups (# subject animals) | 38 |  |  |  |
| Log Likelihood | -1516.4 |  |  |  |
| Akaike Inf. Crit. | 3048.9 |  |  |  |
| Bayesian Inf. Crit | 3080.8 |  |  |  |

**Model formula:** mod\_sm\_quadandnbinom2 = glmmTMB(grooming ~ sex + age + sex\*age + age2 + age2\*sex + (1 | subject), data=all\_data, family="nbinom2", offset=log(totalobs))
**Dispersion parameter for nbinom2 family:** 4.91

**Supporting Information Table 10:** Predicted Values and Simple Effects from Best Grooming Model. The maximum predicted values (before rounding) for each column are in bold.

| Age | Predicted rate for females | Predicted rate for males | Sex difference in rate prediction (M-F) | Rate ratio (M/F) | P-value for the rate ratio |
| --- | --- | --- | --- | --- | --- |
| 0 | 0.0206 | **0.0229** | 0.00228 | 1.11 | 0.492 |
| 1 | 0.0257 | 0.0212 | -0.0045 | 0.825 | 0.153 |
| 2 | 0.0313 | 0.0197 | -0.0116 | 0.63 | 0.000259 |
| 3 | 0.0373 | 0.0185 | -0.0188 | 0.496 | 2.36E-08 |
| 4 | 0.0435 | 0.0175 | -0.026 | 0.402 | 2.31E-12 |
| 5 | 0.0496 | 0.0166 | -0.033 | 0.336 | 1.45E-15 |
| 6 | 0.0553 | 0.0159 | -0.0393 | 0.288 | 7.56E-18 |
| 7 | 0.0603 | 0.0154 | -0.0449 | 0.255 | 2.57E-19 |
| 8 | 0.0642 | 0.0149 | -0.0493 | 0.233 | 4.7E-20 |
| 9 | 0.0669 | 0.0146 | -0.0523 | 0.218 | 4.67E-20 |
| 10 | 0.0682 | 0.0144 | **-0.0538** | 0.211 | 2.89E-19 |
| 11 | **0.068** | 0.0143 | -0.0537 | 0.21 | 1.2E-17 |
| 12 | 0.0663 | 0.0142 | -0.052 | 0.215 | 2.74E-15 |
| 13 | 0.0631 | 0.0143 | -0.0488 | 0.227 | 1.85E-12 |
| 14 | 0.0588 | 0.0145 | -0.0443 | 0.246 | 1.52E-09 |
| 15 | 0.0536 | 0.0148 | -0.0388 | 0.275 | 0.000000652 |
| 16 | 0.0478 | 0.0151 | -0.0326 | 0.317 | 0.0000881 |
| 17 | 0.0416 | 0.0156 | -0.026 | 0.376 | 0.00332 |
| 18 | 0.0354 | 0.0163 | -0.0192 | 0.459 | 0.0405 |
| 19 | 0.0295 | 0.0171 | -0.0125 | 0.578 | 0.205 |

## Notes

https://osf.io/nsxd8/

https://osf.io/jy3w5/

https://osf.io/ybvfg/

### Competing Interest Statement

The authors have declared no competing interest.

